# The serotonin reuptake inhibitor Fluoxetine inhibits SARS-CoV-2

**DOI:** 10.1101/2020.06.14.150490

**Authors:** Melissa Zimniak, Luisa Kirschner, Helen Hilpert, Jürgen Seibel, Jochen Bodem

## Abstract

To circumvent time-consuming clinical trials, testing whether existing drugs are effective inhibitors of SARS-CoV-2, has led to the discovery of Remdesivir. We decided to follow this path and screened approved medications “off-label” against SARS-CoV-2. In these screenings, Fluoxetine inhibited SARS-CoV-2 at a concentration of 0.8µg/ml significantly, and the EC50 was determined with 387ng/ml. Fluoxetine is a racemate consisting of both stereoisomers, while the S-form is the dominant serotonin reuptake inhibitor. We found that both isomers show similar activity on the virus. Fluoxetine treatment resulted in a decrease in viral protein expression. Furthermore, Fluoxetine inhibited neither Rabies virus, human respiratory syncytial virus replication nor the Human Herpesvirus 8 or Herpes simplex virus type 1 gene expression, indicating that it acts virus-specific. We see the role of Fluoxetine in the early treatment of SARS-CoV-2 infected patients of risk groups.

---

Starting in December 2019, SARS-CoV-2 originated in central China became a pandemic thread with more than 7.000.000 cases worldwide and more than 400.000 deaths so far. Despite these high case rates Remdesivir,^1^ the only effective treatment is still not available for most of the patients. At the same time, other promising substances like Lopinavir^2^ and Chloroquine^3^ showed little effect or led to severe adverse side effects. Furthermore, some drugs, such as ribavirin and interferon,^4^ had no significant impact on patient survival rates.

To circumvent time-consuming clinical trials, testing whether existing drugs are effective inhibitors of SARS-CoV-2, has led to the discovery of Remdesivir.^5^ We decided to follow this path and screened approved medications “off-label”^6^ against SARS-CoV-2. In such a trial, we investigated the effect of the serotonin reuptake receptor inhibitors (SRRI) Fluoxetine, Escitalopram, and Paroxetine on viral replication. For that, cytotoxicity and viral replication rates of a patient-derived virus isolate were measured. To investigate cytotoxicity, Vero cells were incubated with the compounds for three days, and cell growth was determined using a Perkin Elmer Ensight reader. The Vero cells were incubated with the compounds at increasing concentrations and subsequently infected with SARS-CoV-2 at an MOI of approx. 0.5. The concentrations were selected near the concentration used for the treatment of depression (e.g., 0.8 µg/ml for Fluoxetine). DMSO was used as solvent control. After three days, viral replication supernatants were collected and viral RNA was extracted using a MagNA Pure 24 system (Roche). Viral replication was quantified by real time RTqPCR with the LightMix Assay SARS-CoV-2 RdRP RTqPCR assay kit (TIB MOLBIOL, Germany) and the RNA Process Control kit (Roche) (Figure 1A). The PCR was pipetted with a pipette robot device to ensure quality (BRAND, Germany). All infections were performed in triplicates and reaped twice. Fluoxetine inhibited SARS-CoV-2 at a concentration of 0.8µg/ml significantly, and the EC50 was determined with 387ng/ml (Figure 1A). However, it is unlikely that the direct inhibition of the serotonin reuptake transporter is responsible for this suppression since neither Paroxetine nor Escitalopram interfere with viral replication. Furthermore, since most of the cellular off-target factors of the different compounds are similar, Fluoxetine presumably targets the virus directly. Fluoxetine is a racemate consisting of both stereoisomers, while the *S*-form is the dominant SSRI.^7^ Thus, we analyzed which stereoisomer is inhibiting SARS-CoV-2 replication. We found that both isomers show similar activity on the virus (Figure 1A) underlining that the antiviral effect is unrelated to the serotonin reuptake receptor. To get further insights into the mechanism of inhibition, we sought to visualize viral protein expression by immunofluorescence with a patient-derived antiserum. Fluoxetine treatment resulted in a decrease in protein expression, showing that Fluoxetine acts upstream of gene expression (Figure 1B). To further analyze the specificity of Fluoxetine, the inhibition of other viruses was determined. Fluoxetine inhibited neither Rabies virus, human respiratory syncytial virus (RSV) replication nor the Human Herpesvirus 8 or Herpes simplex virus type 1 gene expression, indicating that it acts virus-specific (Figure 1C).

**Figure 1:**
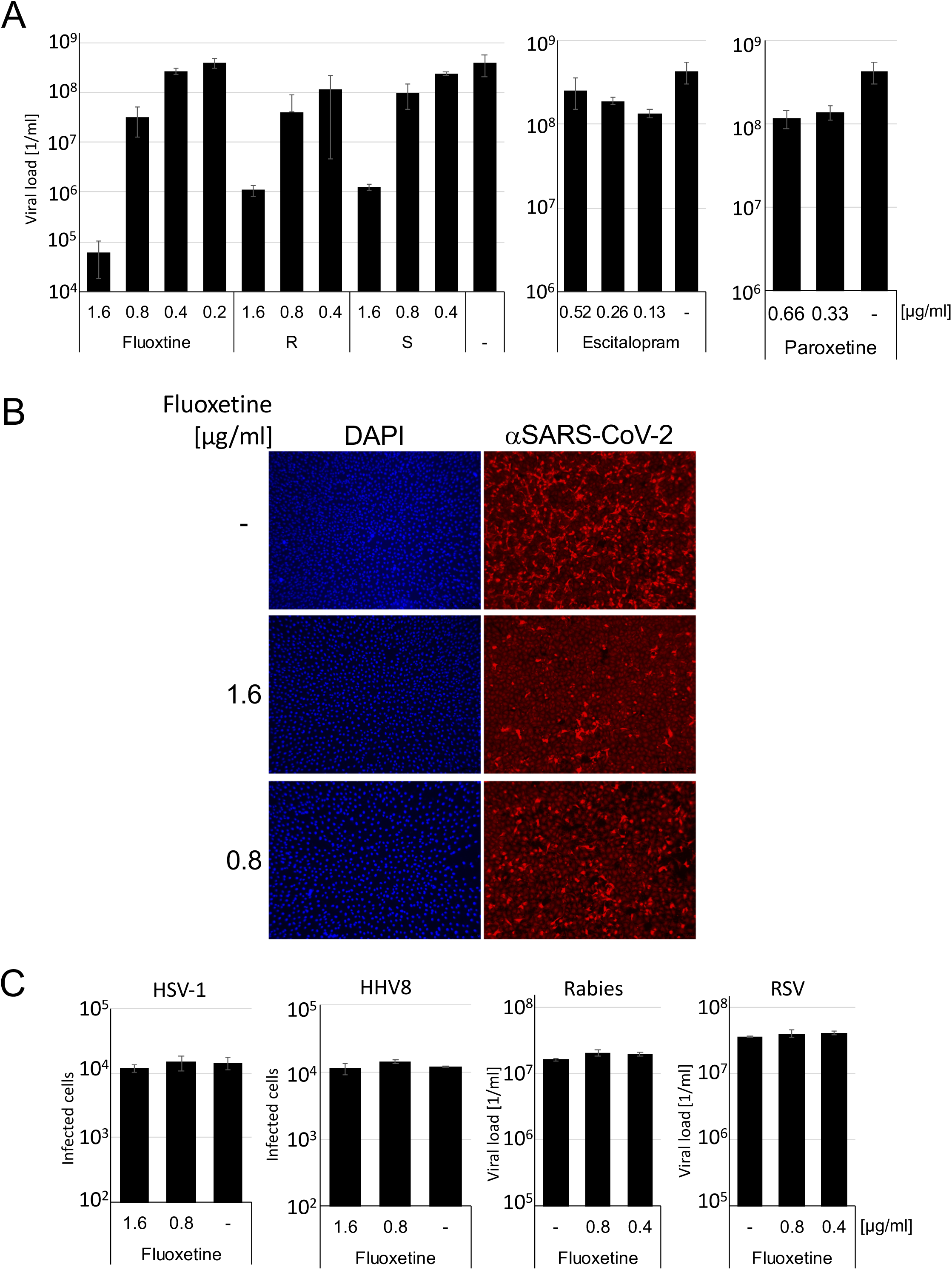
Fluoxetine inhibits SARS-CoV-2 replication. **A**. Vero cells were incubated with the compounds and subsequently infected with SARS-CoV-2 (S: S-stereoisomer; R: R-stereoisomer). Cellular supernatants were collected three days after infection, and viral titers were determined with RTqPCR. **B**. Vero cells were infected with SARS-CoV-2 for 72h and viral proteins were detected with a SARS-CoV-2 specific antiserum (1:100) and a TexasRed-labeled donkey anti-human antibody (1:500, Dianova). The nuclei were stained with DAPI. **C**. BHK21 and HepG2 cells were infected with either *gfp*-encoding HSV-1, HHV8, with a Rabies vaccine strain or with a patient derived RSV. Viral titers were determined by RTqPCR (RSV, Rabies) or infected *gfp*-expressing cells were counted with an Ensight device (PerkinElmer).

Fluoxetine was introduced in clinics during the seventies and is a well-studied drug since it has been used in humans for almost four decades. Furthermore, the patent of Fluoxetine has long expired. Thus, it is available from different companies, and relatively cheap. We see the role of Fluoxetine in the early treatment of SARS-CoV-2 infected patients of risk groups.

## Acknowledgement

We would like to thank Novartis Germany and Jörg Vogel for support.

